# GENE THERAPY PREVENTS HEPATIC MITOCHONDRIAL DYSFUNCTION IN MURINE DEOXYGUANOSINE KINASE DEFICIENCY

**DOI:** 10.1101/2024.05.10.593325

**Authors:** Nandaki Keshavan, Miriam Greenwood, Helen Prunty, Juan Antinao Diaz, Riccardo Privolizzi, John Counsell, Anna Karlsson, Simon Waddington, Rajvinder Karda, Shamima Rahman

## Abstract

Primary mitochondrial disorders are an uncommon cause of neonatal hepatic failure. Biallelic pathogenic variants of the gene encoding the mitochondrial localising enzyme deoxyguanosine kinase (DGUOK) cause hepatocerebral mitochondrial DNA depletion syndrome leading to acute neonatal liver failure and early mortality. There are currently no effective disease-modifying therapies. In this study, we developed an adeno-associated virus 9 (AAV9) gene therapy approach to treat a mouse model of DGUOK deficiency that recapitulates human disease. We delivered AAV9-*hDGUOK* gene therapy intravenously to newborn *Dguok* knock-out mice and showed that liver dysfunction was prevented in a dose dependent manner. Unexpectedly for neonatal delivery, durable and long-lasting liver transduction and RNA expression were demonstrated. Liver mitochondrial DNA depletion, deficiencies of oxidative phosphorylation complexes I, III and IV and liver transaminitis and survival were ameliorated in a dose-dependent manner.

## Introduction

Mitochondrial DNA depletion (mtDNA) syndromes (MDDS), comprise a heterogenous subgroup of severe primary mitochondrial disorders caused by mutations in nuclear genes encoding key proteins involved in mtDNA replication or maintenance of mitochondrial deoxyribonucleotide triphosphate (dNTP) pools^1^. Nucleotides incorporated into mtDNA are either synthesized *de novo* or salvaged from the cytosol via a series of biochemical steps known as the mitochondrial salvage pathway. Defects in any of ∼14 genes result in mtDNA depletion and therefore loss of the mtDNA-encoded subunits of oxidative phosphorylation (OXPHOS) enzymes, lead to severe energy deficiency frequently presenting as debilitating infantile-onset disease with a high mortality^2^.

Biallelic pathogenic variants in *DGUOK* encoding deoxyguanosine kinase, a component of the mitochondrial nucleoside salvage pathway, account for up to 20% of MDDS^3^. DGUOK catalyses the intramitochondrial phosphorylation of dG and dA to dGMP and dAMP respectively^4^. Loss of DGUOK function results in imbalanced nucleotide pools, nucleotide misincorporation and mtDNA depletion^5^. Neonates and infants with DGUOK deficiency typically present with severe acute liver failure which correlates with early mortality. Current management is only supportive. Advances in AAV-based gene therapy technology and subsequent clinical trial successes have led to market approval for a few genetic disorders but this is not yet the case for primary mitochondrial diseases.

There is a mouse model of the DGUOK deficiency, which closely recapitulates the human disease phenotype. The model generated by disruption of exon 2 of *Dguok* using Cre/lox homologous recombination, demonstrates weight loss and chronic liver disease. In this study, we aimed to develop an AAV9-based gene therapy approach to prevent liver mitochondrial dysfunction in the *Dguok* KO mouse model. We showed that AAV9 mediates efficient lasting liver transduction enabling rescue of mtDNA depletion, OXPHOS abnormalities, transaminitis and a dose-dependent improvement in survival in KO mice.

## Results

### Dguok KO mice recapitulate the human disease phenotype

KO mice demonstrated liver mtDNA depletion from birth with mtDNA levels of ∼29% of WT/HET controls. Liver mtDNA copy number decreased further to <5% of WT/HET controls by 3 months’ age and remained at this level thereafter (**Figure 1A**).

**Figure 1:**
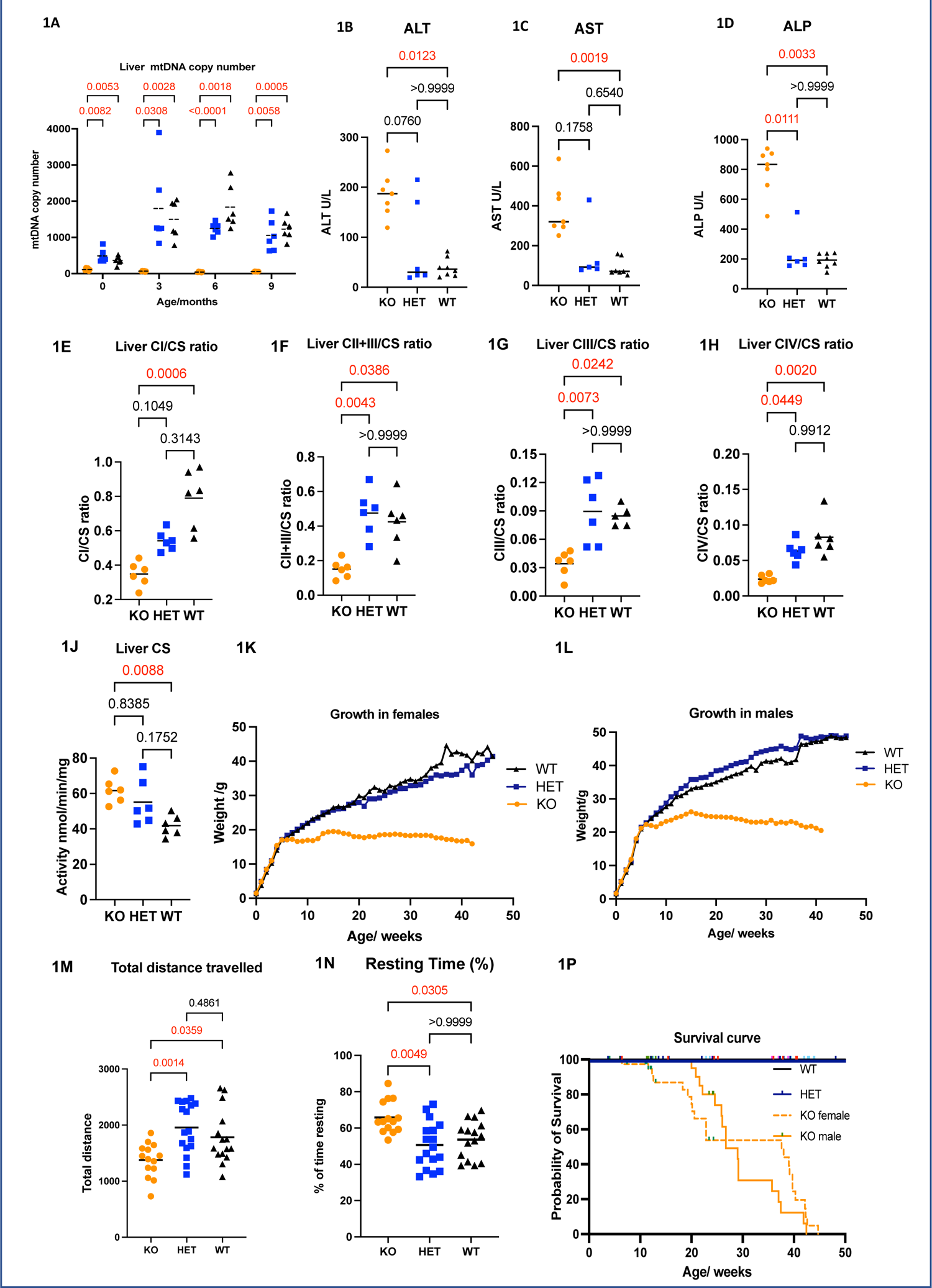
Phenotyping of murine *Dguok* KO model **1A:** Liver mtDNA quantitation. KOs showed significant liver mtDNA depletion compared to WTs. **1B-1D:** Liver function tests. KOs showed increased blood ALT, AST and ALP levels. **1E-1J:** Multiple OXPHOS abnormalities demonstrated in liver. **1K-1L:** Female and male KOs showed lower body weights compared to WTs from 6 weeks (indicated by the arrows). **1M-1N:** Open field testing. KOs show reduced total distance travelled and increased resting time compared to WTs. **1P:** Survival curve of *Dguok* knock-out mice as determined by reaching the humane endpoint. Upward ticks indicate censored subjects. Sample size: 23 WTs, 53 HETs and 76 KOs. Statistics: Graphs show means, p values are indicated above the brackets. Kruskal-Wallis test used for all analyses except survival for which Mantel-Cox test was used.

Elevations of ALT, AST, ALP (**Figures 1B-D**), the amino acids threonine, glycine, arginine, methionine (**Figures S1A-D**) elevated ammonia **(Figure S1E**) were seen in KO mice. OXPHOS activities in tissue homogenates from mice at 9 months’ age revealed multiple OXPHOS deficiencies in liver of KO mice compared to WTs. Complex I, II+III, III and IV activities were deficient, together with significant elevation of citrate (CS) synthase activity reflecting a compensatory increase in mitochondrial mass (**Supplementary Table 1** and **Figures 1E-J**).

In brain, KOs had mtDNA copy numbers of ∼50% of WT/HET levels at birth and at 9 months this was ∼40% of WT/HETs (**Figure S1F)**. From a biochemical perspective, brain was affected more mildly showing only isolated complex IV deficiency (**Figure S1L**). Increased GFAP expression was observed throughout the brain of some KO mice, but this finding was variable. Statistically significant increases in GFAP expression in KOs compared to WTs were observed in olfactory nucleus, cortex, striatum and medulla oblongata (**Figure S2**).

MtDNA depletion was also observed in skeletal muscle, heart and spleen and a deficiency of mtDNA copy number was seen in kidney (**Figure S1G-K**). Complex I deficiency was seen in skeletal muscle (**Figure S1M**). No OXPHOS abnormalities were observed in heart (**Table 1**).

Growth velocity of KOs decreased from 6 weeks of age in both sexes. KOs reached a maximum weight (19g in females and 25g in males) at about 16 weeks, after which their weights plateaued and eventually declined (**Figures 1K, 1L).** On open field testing, KOs demonstrated significantly lower total distance travelled and increased resting time, as shown in **Figures 1M, 1N.** There was no significant difference in grip strength in KO mice **(Figure S1N).**

The humane endpoint was defined as 15% weight loss from the highest measured weight. All KO mice reached this endpoint by 42 weeks whereas all WT and HET mice survived (**Figure 1P**). Female and male KOs had median survival of 37.5 and 26.7 weeks respectively. This difference was statistically significant (Mantel-cox test, p<0.0001)

### Gene transfer ameliorates liver disease in *Dguok* KO mice

Of all organs, the highest transduction was seen in liver. Vector copy number (VCN) was significantly higher in the 8×10^14^vg/kg KO group (mean VCN 5.03) than the 8×10^13^ vg/kg KO group (mean VCN 1.57, p=0.0116), indicating a dose response (**Figure 2A**). VCN data in skeletal muscle and heart showed lower transduction than liver (**Figure S4A, S5A).** In liver, *hDGUOK* expression normalised to *mGapdh* in injected KOs exceeded endogenous *mDguok* expression at both doses (8×10^13^vg/kg KOs: mean *hDGUOK* expression 1.72, 8×10^14^vg/kg KOs: mean *hDGUOK* expression 1.81, uninjected WT *mDguok* expression: 0.018, corresponding to fold changes of 95 and 100 respectively). There was also greater *hDGUOK* RNA expression in injected KOs compared to dose-matched injected WTs. WT injected mice had *mDguok* levels that were similar to control uninjected WTs i.e. no downregulation of endogenous gene expression (**Figure 2B**).

**Figure 2:**
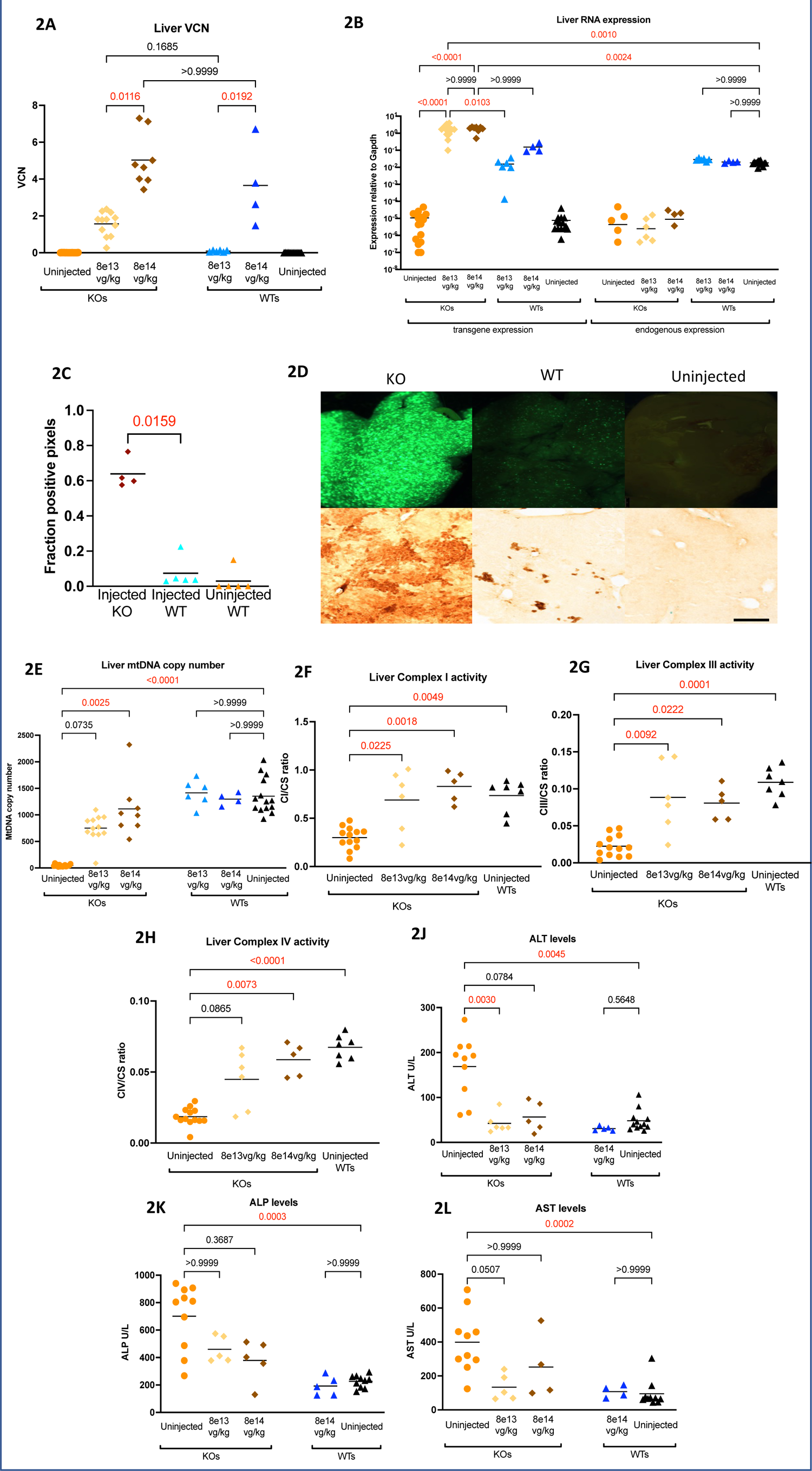
Efficacy of liver-directed gene therapy. **2A:** Liver VCN. **2B:** Liver RNA expression. **2C** GFP Quantitation was performed on images acquired at x40 magnification, statistical analysis: Mann Whitney test comparing the two injected groups only, p value shown above the bracket. Scale bar: 500µm. Sample sizes: injected KO: 4, injected WT: 5, uninjected WT: 5. **2D** Stereoscopic imaging, anti-GFP immunohistochemistry in liver in long-term biodistribution study. Stereoscopic imaging (left column): Exposure was identical for the three groups (700ms). Immunohistochemical images (right column): images were acquired at ×10 magnification. **2E:** Liver mtDNA copy number. **5F-5H:** Liver complex I, III and IV activity. **5J-5L:** Liver function tests. Statistics: Kruskal-Wallis test.

In biodistribution studies, quantification of GFP expression by immunohistochemistry revealed a >6-fold increase in long-term GFP expression in injected KO liver compared to injected WTs (p=0.0159). The pattern of GFP positivity was non-uniform with islands of positive cell clusters interspersed between clusters of negative cells (**Figure 2C-D**). In skeletal muscle and cardiac tissue, RNA expression showed dose-responsiveness which was supranormal at the 8×10^14^vg/kg dose (**Figure S4B, S5B).**

Rescue of mtDNA copy number was observed in liver from a baseline of 3% in uninjected KOs, to 55% for the 8×10^13^vg/kg KO group and 82% for the 8×10^14^vg/kg KO group (**Figure 2E**). No mtDNA abnormality was observed in injected WTs. Dose-dependent improvement in mtDNA copy number was also demonstrated in skeletal muscle (**Figure S4C).** No significant improvement in heart mtDNA copy number was observed **(Figure S5C).** Rescue of complex I, III and IV activities was observed in liver in a dose-dependent manner (**Figure 2F-H**) and of complex I in skeletal muscle (**Figure S4D).** Complete rescue of ALT levels was seen at both doses (8×10^13^ vg/kg and 8×10^14^ vg/kg) in injected KOs, together with partial improvements in AST and ALP. Liver function remained normal in injected WTs (**Figure 2J-L**).

### Gene transfer does not improve brain abnormalities in *Dguok* KO mice

In contrast to the liver, in the brain transduction was poor. VCNs were lower (8×10^14^vg/kg group mean VCN: 0.09, 8×10^13^ vg/kg group mean VCN: 0.05) compared to liver. No significant difference was observed in mean VCNs in injected KOs and WTs (∼0.11 in both groups) **(Figure S3A)**. In brain, injected KOs also showed comparably lower *hDGUOK* RNA expression than in liver and *hDGUOK* expression was not significantly different to WT endogenous *mDguok* expression at either dose **(Figure S3B**). In biodistribution studies, GFP expression, though present, was generally low (data not shown). There was persistent mtDNA depletion (22% in uninjected KOs, 26% in the 8×10^13^vg/kg KO group and 27% in the 8×10^14^vg/kg KO group, **Figure S3C)** associated with persistent complex IV deficiency at both doses (Figure S3D).

### Growth, locomotor and survival outcomes

We observed minimal improvement of growth in KO injected female mice and no improvement in males after gene therapy. Growth was significantly impaired in WT injected mice at the 8×10^14^ vg/kg dose implying there was a negative impact on growth. No significant growth abnormality was seen in WTs injected at the 8×10^13^ vg/kg (**Figure 3A-B**). Locomotor abnormalities were partially ameliorated in AAV9 injected KOs with respect to both total distance travelled and percentage resting time at both doses compared to uninjected WTs (**Figure 3C-D**). Survival was ameliorated in a dose-dependent manner in injected KO mice (baseline median survival 177 days, 8×10^13^vg/kg KO group 261 days, 8×10^14^vg/kg KO group 100% survival until end of follow up at 42 weeks). For the 8×10^13^vg/kg KO group this was not significantly different to uninjected KOs (Mantel-cox test, p=0.09). WT mice injected at both 8×10^13^vg/kg and 8×10^14^vg/kg had 100% probability of survival in long-term follow up. High-dose AAV9 (8×10^15^ vg/kg) caused early mortality in both WTs and KO mice with a median survival of 20 days, suggesting toxicity (**Figure 3E-F**).

**Figure 3.**
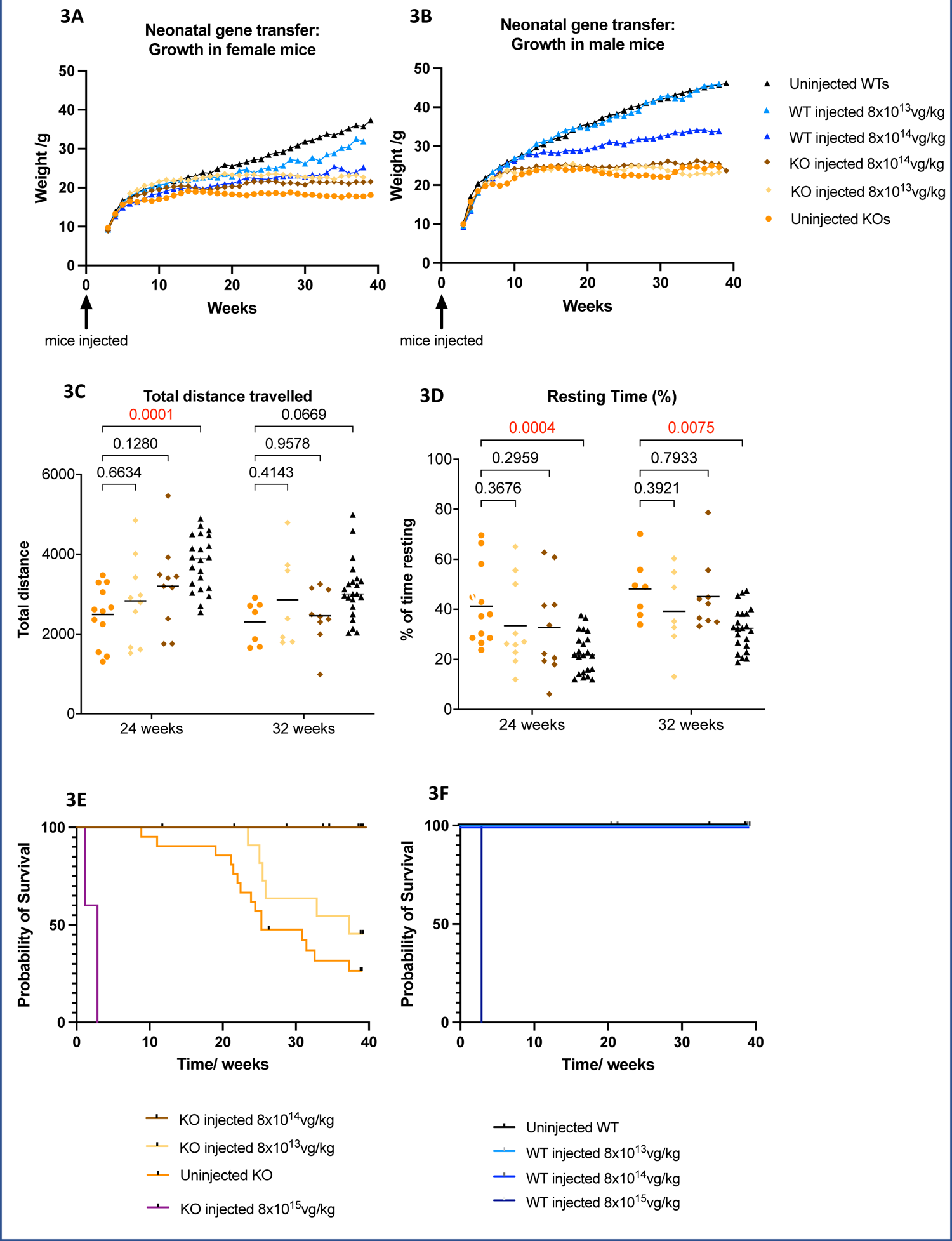
Growth, behavioural and survival outcomes following neonatal gene transfer. **3A** and **3B:** Growth outcomes in female and male mice following neonatal gene transfer. **3C and 3D:** Total distance travelled and percentage resting time in injected mice. **3E-F:** Survival in injected KO and WT mice expressed as a probability vs time. Statistics: Graphs show means, p values are indicated above the brackets. Kruskal-Wallis test used for all analyses except survival for which Mantel-Cox test was used.

## Discussion

Infantile-onset DGUOK deficiency is characterised by progressive liver failure including conjugated hyperbilirubinaemia, coagulopathy, transaminitis, tyrosinaemia, and histological evidence of cholestasis, microsteatosis, fibrosis, haemochromatosis, cirrhosis, necrosis, portal hypertension, hepatocellular carcinoma.^6-13^ Management of DGUOK deficiency is supportive, including management of hypoglycaemia, cholestasis, and the complications of liver failure.^14^ Liver transplants in 14 patients showed unsatisfactory results: 1-year and 5-year post-transplant survival were 64% and 35% respectively which is lower than the average for liver transplantation for all causes.^15, 16^ Four patients had severe neurological progression despite liver transplantation, including one who had no apparent neurological disease prior to liver transplant. These data and the shortage of suitable organ donors underscore a need for novel effective therapies. AAV-based gene therapy has been used to treat several mouse models of mitochondrial disorders^17–23^ but so far none of yet been licensed. Success in treating DGUOK deficiency with gene therapy may widen the scope for treating other genetic causes of mitochondrial DNA depletion syndrome.

Liver dysfunction is a core feature of DGUOK deficiency. It has previously been shown in other studies that liver transgene expression following neonatal gene transfer in mice is short-lived due to a dilutional effect as the liver grows^24, 25^. In this work, long-term biodistribution studies demonstrated KO mice had significantly higher GFP expression than WTs in liver. Islands of GFP positive cells were seen in KO livers and the distribution of overall positivity appeared to be higher than that observed in the short term, contrary to expectations. These intriguing results raised the possibility that the AAV vector might be integrating into the host cell genome. AAV integration is a well-known phenomenon. Recent studies have suggested that the frequency of recombinant AAV integration may be higher than previously recognised, at a frequency of 1-3% in liver^26^.However, integration alone would not explain why only KO and not WT mouse livers showed high GFP expression. The possibility that underlying mitochondrial dysfunction in KO mice may influence transgene expression needs to be considered. One explanation could be that mitochondrial dysfunction leads to greater cell turnover in KO mice that causes cell death. Transduced KO hepatocytes would be cured of their mitochondrial dysfunction and therefore could be conferred a survival advantage whilst non-transduced cells succumb to disease. Over time, cured cells could repopulate the liver resulting in high VCN and GFP expression. Similar observations have been seen in another metabolic liver disease, fumarylacetoacetate hydrolase (FAH) deficiency^27^. Another possibility is that higher GFP expression in KO mice may not lie at the level of cellular transduction, but more distally at the level of RNA or protein expression. Indeed we observed significantly higher transgene RNA expression in injected KOs compared to WTs delivered the same AAV dose.

Furthermore, this observation has dose-implications since the dose needed to achieve high long-term transgene expression could be lower than anticipated, which is a major positive consideration given that toxicity is well recognised at high AAV doses. Other unanswered questions include whether other liver mitochondrial diseases may exhibit the same phenomenon seen in the DGUOK mouse model, and whether the same would be seen in humans. If this were true then AAV9-based gene therapy could be a preferred treatment for mitochondrial diseases involving the liver.

The clinical definition of MDDS is <30% of healthy controls and the mouse model of DGUOK deficiency faithfully demonstrates liver mtDNA content <5% of WT levels. Efficacy of liver-directed neonatal gene therapy was evaluated at AAV9 doses of 8×10^13^ vg/kg and 8×10^14^ vg/kg. VCN studies showed excellent dose-dependent transduction of liver in injected KO mice (1.57 and 5.03/cell for the 8×10^13^ vg/kg and 8×10^14^ vg/kg dose groups respectively) corresponding to 79.2 and 99.3% cellular transduction. Mean *hDGUOK* RNA expression in injected KOs was ∼ 100 fold higher than mean endogenous WT *mDguok* levels, in both the 8×10^13^ vg/kg and 8×10^14^ vg/kg groups suggesting saturation of RNA expression at this dose range. Therapeutic efficacy was observed in liver at both 8×10^13^ vg/kg and 8×10^14^ vg/kg, restoring liver mtDNA content to 55% and 82% of WT levels respectively. For the 8×10^14^ vg/kg group there was no statistical difference compared to WTs implying complete rescue. For the 8×10^13^ vg/kg group, there was partial rescue despite the high *hDGUOK* RNA expression as discussed above. Human and mouse *Dguok* cDNA have 80.6% identity corresponding to 75.1% identity at an amino acid level, however their 39 amino acid long mitochondrial targeting peptides which enable uptake of DGUOK protein into mitochondria are quite dissimilar (∼48% identity). Therefore, it is possible that even if hDGUOK protein were overexpressed proportionately to *hDGUOK* RNA, species differences in mitochondrial targeting could potentially lead to a decrease in hDGUOK localisation to mitochondria. Despite these species differences, we see a significant amelioration of mtDNA depletion to well above the clinical definition (30% of WT controls) at both doses. Rescue of mtDNA copy number in the 8×10^13^ vg/kg and 8×10^14^ vg/kg groups helps to answer the question of what proportion of transduced hepatocytes is required to ensure sufficient rescue of mtDNA copy number to >30% of WT levels. Our results suggest that 35.1% transduced cells are needed to achieve a mtDNA content of 30% (data not shown). Having demonstrated amelioration of mtDNA copy number in liver, we then proceeded to investigate whether OXPHOS abnormalities and transaminitis were rescued. We observed an improvement in complex I, III and IV activities at both doses. The extent of rescue for the lower dose (complex I to 93% of WT levels, complex III to 81% of WT levels, complex IV to 66% of WT levels) is particularly noteworthy because it implied that a mean mtDNA copy number of 55% was sufficient to achieve significant rescue of liver mitochondrial dysfunction and subsequent amelioration of liver transaminases.

Patients with infantile-onset DGUOK deficiency also develop neurological disease including hypotonia, developmental delay, ptosis, rotatory nystagmus, and seizures and the DGUOK KO mouse model recapitulates the brain phenotype by demonstrating decreased mtDNA levels and complex IV deficiency. Although targeting of the liver through IV injections was the primary aim of this study, we also assessed brain targeting via this route. Long-term biodistribution studies demonstrated poor brain transduction in both WTs and KOs. In the gene therapy experiments, VCNs at either dose (8×10^13^ vg/kg or 8×10^14^ vg/kg) were low (0.05 and 0.09/cell respectively) implying low transduction. It was therefore unsurprising to see no significant improvement in mean mtDNA copy number or complex IV activity.

Improved gene transfer to the brain is needed by employing alternative strategies or intracranial routes of delivery. In another mouse model of MDDS caused by *TWNK* deficiency which is associated with encephalopathic MDDS in humans, mtDNA depletion in glia specifically led to astrogliosis, analogous to our observations in the DGUOK deficient mouse model^28^. These data suggest that adequate glial transduction is essential to rescue neurological involvement in this model.

Although we saw complete correction of liver disease, we did not observe rescue of growth. These data suggest that involvement of other organs could be contribute to poor weight gain, for example the brain, kidney, skeletal muscle, gastrointestinal or endocrine systems. Further work is needed to interrogate this. We also assessed locomotion in injected mice. The main abnormalities found in the KO strain were increased % resting time and reduced total distance travelled. Neonatal gene transfer appeared to partially improve total distance travelled and % resting time. These data suggest that locomotor abnormalities in KOs may also have a multisystemic aetiology.

Survival analyses showed a dose-dependent amelioration of survival, with complete rescue of survival at 8×10^14^ vg/kg and non-significant improvement at the 8×10^13^vg/kg dose. As liver transaminitis was significantly ameliorated at both doses these data suggest that the involvement of other organs in the model could explain these findings. One possibility is skeletal muscle involvement since we observed a dose-dependent improvement in skeletal muscle mtDNA content but it should be noted that myopathic disease is not a prominent clinical feature of infantile-onset DGUOK deficiency. These data imply that reduced survival in this model is also likely to be due to multisystemic aetiology.

We sought to clearly define an upper limit for dosing our AAV vector by using a dose of 8×10^15^vg/kg in the highest dose group. This caused toxicity in both KO and WT mice resulting in early death. At the 8×10^14^vg/kg or 8×10^13^vg/kg doses, there was no increase in mortality seen in injected WT mice, however, there was a decrease in growth at 8×10^14^vg/kg. However, for the lower 8×10^13^vg/kg dose group, growth was normal. These data suggest a dose-dependent toxic effect on growth in injected WTs. The mechanisms underlying this toxicity are unclear since mtDNA copy number in liver, brain or skeletal muscle and blood liver function tests were normal in injected WTs. Nevertheless, it is clear that IV doses ≥ 8×10^14^vg/kg are not justified. It is also important to note that from a clinical perspective, doses above 1×10^14^vg/kg have been associated with toxicity^29^.

We aimed to also define the minimum efficacious dose needed to rescue liver disease. So far both the 8×10^13^vg/kg and the 8×10^14^vg/kg doses were able to rescue liver mtDNA content to >30% of WT controls, and completely normalised ALT levels. Further IV dose de-escalations would be needed to ascertain minimum efficacious dose for liver-directed gene therapy in this model.

Considering the dosing requirements of the two main organs involved in DGUOK deficiency (the liver and brain), it is clear that achieving efficacious targeting of both organs using a single neonatal IV dose of this AAV9 construct will be challenging, since 8×10^14^vg/kg was unable to transduce the brain sufficiently and higher doses appear toxic. Alternative vectors or intracranial routes of delivery need to be considered to achieve efficacy and safety.

## Materials and Methods

### Cloning of plasmids

Two payload plasmids were generated, pAAV-CAG-intron-*hDGUOK-*T2A-eGFP-WPRE-bGH-pA and pAAV-CAG-intron-*hDGUOK*-WPRE-bGH-pA, for biodistributional and gene therapy studies respectively. First, pAAV-CMV-GFP-WPRE-bGH-pA (University of Pennsylvania) was linearised by PCR (primer sequences available on request). Gene blocks containing *hDGUOK-T2A and hDGUOK* were obtained from Integrated DNA technologies (IDT, Leuven, Belgium). The CAG promoter and intron were restriction digested from existing plasmids. Ligation of the various components was undertaken using an In-Fusion cloning kit (Takara Bio-Europe, Paris, France). The ligation reaction was transformed into Stellar competent cells as per manufacturer protocols. Cells were incubated on LB/agar plates overnight and clones selected. The DNA sequences obtained from clones were confirmed using Sanger Sequencing.

### AAV9 vector production and titration

DNA was amplified for AAV production using an Invitrogen maxiprep kit. Helper and AAV9 plasmids (Harvard University and University of Pennsylvania respectively) were used in AAV production using a triple transfection approach in HEK293T cells, followed by HPLC purification (AKTA prime) and vector concentration as described previously^30^. Following DNAse treatment, the vector was titrated via qPCR using Luna SYBR green reagents (New England Biolabs) as per manufacturer recommendations. Primer sequences targeting *hDGUOK* were used for titration and are available on request. qPCR standards were made up using gene blocks as above.

### Animal experiments

#### Husbandry

Experimental animals were maintained at an experimental animal facility in adherence with Animal Research: Reporting of In Vivo Experiments (ARRIVE) guidelines and to UK Home Office regulations. Experiments were approved by UCL biological services. Mice were housed in individually ventilated cages, subject to day/ night light cycles, provided with drinking water, standard laboratory rodent chow and nesting materials.

#### Breeding

Adult *Dguok*^+/-^ mice were maintained on an albino C57BL/6N background for breeding of *Dguok*^-/-^ mice (KOs) which were identified by genotyping as described previously^31^.

#### Behavioural testing

Open field testing was carried out using Harvard Panlab equipment (Barcelona, Spain) in 25cmx25cm square arenas. Recordings of 10 minute duration were taken in moderate lighting. Data were analysed using SMART v3 software. For grip strength testing animals were placed onto a 1cm x 1cm metal grid and then gently inverted over a large transparent plastic box. The time taken to fall in seconds was recorded (average of three attempts taken as the final measurement).

#### Intravenous injections

Injections were performed within the first 48 hours of life via superficial temporal vein using a 33-gauge Hamilton needle. 20 µl of vector was administered per pup. Investigators undertaking gene therapy experiments were blinded as to which animals received injections and animals were randomised to treatment groups. Both male and female mice were used. In initial biodistribution studies, WTs were injected at birth with AAV9-CAG-*hDGUOK*-GFP at 3×10^13^vg/kg or 3×10^14^vg/kg and followed up for 6 weeks. Tissues were collected and stained for GFP. We assessed long-term GFP expression in both WTs and KOs injected with AAV9-CAG-*hDGUOK*-GFP at 3×10^14^vg/kg only. and followed up to 9 months or the humane endpoint, whichever was sooner, to determine longevity of transgene expression by stereoscopic microscopy, anti-GFP immuno-histochemistry and vector copy number (VCN) studies.

Gene therapy studies were undertaken in neonatal KO mice, using the gene therapy vector, AAV9-CAG-*hDGUOK*. WT littermates were also injected for toxicity studies. The doses used were 8×10^13^vg/kg, 8×10^14^vg/kg, 8×10^15^vg/kg. Animals were followed up until 9 months or the humane endpoint, whichever was first.

#### Collection and processing of tissues

For blood sampling and immunohistochemistry studies, animals were anaesthetised using isoflurane. Blood was collected via intracardiac route and whole-body perfusion undertaken with 1xPBS. Serum samples were obtained after centrifugation of clotted blood at 13500 rpm for 6 minutes. Tissues were fixed in 4% paraformaldehyde for 48 hours, transferred to 30% sucrose, then sectioned to 40nm sections using a microtome (Epredia) and stored at 4 °C in TBSAF. Tissues for OXPHOS studies were obtained by cervical dislocation without anaesthesia and snap frozen in dry ice. Homogenates were prepared for OXPHOS studies as described previously ^32^. DNA and RNA were extracted using Qiagen DNeasy and Invitrogen RNA extraction kits as per manufacturer protocols.

### ddPCR

Tissue VCN and mtDNA copy numbers were determined using ddPCR. The targets used were *hDGUOK* and *Mt-Nd1*, with *Rpp30* as the reference (Bio-rad assay catalogue numbers 10042958 and 10042961 respectively). Primers and probe sequences for *hDGUOK* are available on request. Samples were first prepared by restriction enzyme digestion with *HaeIII* (NEB) for 1 hour at 37 °C followed by ddPCR using a Bio-rad Auto DG droplet generator. Thermocycler settings were: initial activation 95 °C 10mins, denaturation 94 °C 30s, annealing extension 55.8 °C 1 min, cycles 40, deactivation 98 °C 10mins, 4 °C hold. Samples were then read by the Bio-rad droplet reader and analysed using QuantaSoft software v1.7 regulatory edition.

### qPCR

qPCR was undertaken for RNA expression studies, utilising *hDGUOK* and *mDguok* targets, with *mGapdh* as the reference and NEB Luna mastermix for probes. Final concentrations were 450nm for target primers and probes and 250nM for *mGapdh* primers and probe. Primer and probe sequences are available on request. qPCR standards were made up using Gene blocks obtained from IDT for all three targets and were run alongside each qPCR plate. Samples were run in triplicate. Thermocycler settings: initial activation 50 °C 2 mins, initial denaturation 95 °C 10mins, denaturation 95 °C 15s, annealing/extension 60 °C 1 min, cycles 40, hold 4 °C. Final expression data were expressed as a ratio to *mGapdh* expression.

### OXPHOS studies

OXPHOS studies (complexes I, II+III, III, IV and citrate synthase as a reference enzyme) were undertaken as previously described^32–36^.

### Immunohistochemistry and microscopy

Immunohistochemical staining for GFAP-positive astrocytes and CD68-positive microglia was undertaken to investigate the possibility of astrogliosis and microgliosis in baseline phenotyping. Anti-GFP immunohistochemistry was used to assess biodistribution. Free-floating staining of brain and visceral organs was undertaken as previously described^37^. Primary antibodies used: rabbit anti-GFP for GFP staining (Abcam, dilution 1:10000), mouse anti-GFAP for GFAP staining (Merck, 1:1000) and rat anti-CD68 for CD68 staining (Bio-rad, 1:100). Secondary antibodies used: goat anti-rabbit for GFP staining (1:1000), goat anti-mouse for GFAP staining (1:1000) and rabbit anti-rat for CD68 staining (1:1000) (all from Vector laboratories). A 3,3’-diaminobenzidine reporter was used.

### Blood tests

An NX600 Dri-chem analyser (Fujifilm) was used to measure blood liver function tests (ALT, AST, ALP, albumin, bilirubin) and blood glucose from serum as per manufacturer recommendations. Ammonia was measured from whole blood using a N×10N analyser (Fujifilm) as per manufacturer recommendations. Amino acids were analysed as phenylisothiocyanate derivatives by reverse-phase HPLC chromatography using an ODS-bonded silica column (Waters WAT010950) and UV detection at 254 nm, based on a previously reported methods^38, 39^.

### Statistical analysis

Graphpad prism software v9 was used for statistical analysis. p<0.05 was considered statistically significant. For all analyses (except for survival) non-parametric tests (Kruskal-Wallis test and Mann-Whitney test) were used to compare groups rather than parametric equivalents due to data not being normally distributed. For survival analyses, Mantel-cox test was used.

## Data Availability

Data sets will be made available on request

## Acknowledgements

NK received an Action Medical Research Clinical Research Training Fellowship award GN2682 to undertake this work. SR acknowledges grant funding from Great Ormond Street Hospital Children’s Charity, the Lily Foundation and the National Institute of Health Research (NIHR) Great Ormond Street Hospital Biomedical Research Centre. SW received support from MRC grant MR/T016809/1, Action Medical Research grant GN2647, Action Medical Research-Borne grant GN2984 and support from Wellbeing of Women. RK received support from LifeArc grant P2020-0008 and Great Ormond Street Hospital Children’s Charity grant V4720. JAD received support from LifeArc grant P2020-0008. RP was funded through a UCL School of Life and Medical Science Impact PhD Studentship. AK was funded by Karolinska Institute grant 15–0953; Swedish Cancer Society grant CAN 2016/1342-1345, Swedish Research Council grant K2014-66X12162-18-3. The views expressed are those of the authors and not necessarily those of the NHS, the NIHR, or the Department of Health.

## Authors contributions

Conceptualisation: NK, RK, SW, SR, JC

Experiments: NK, MG, HP, RK, SW, JD, RP

Drafting Manuscript and Figures: NK

Review of manuscript: all authors

## Supplementary figure legends

**Figure S1.**
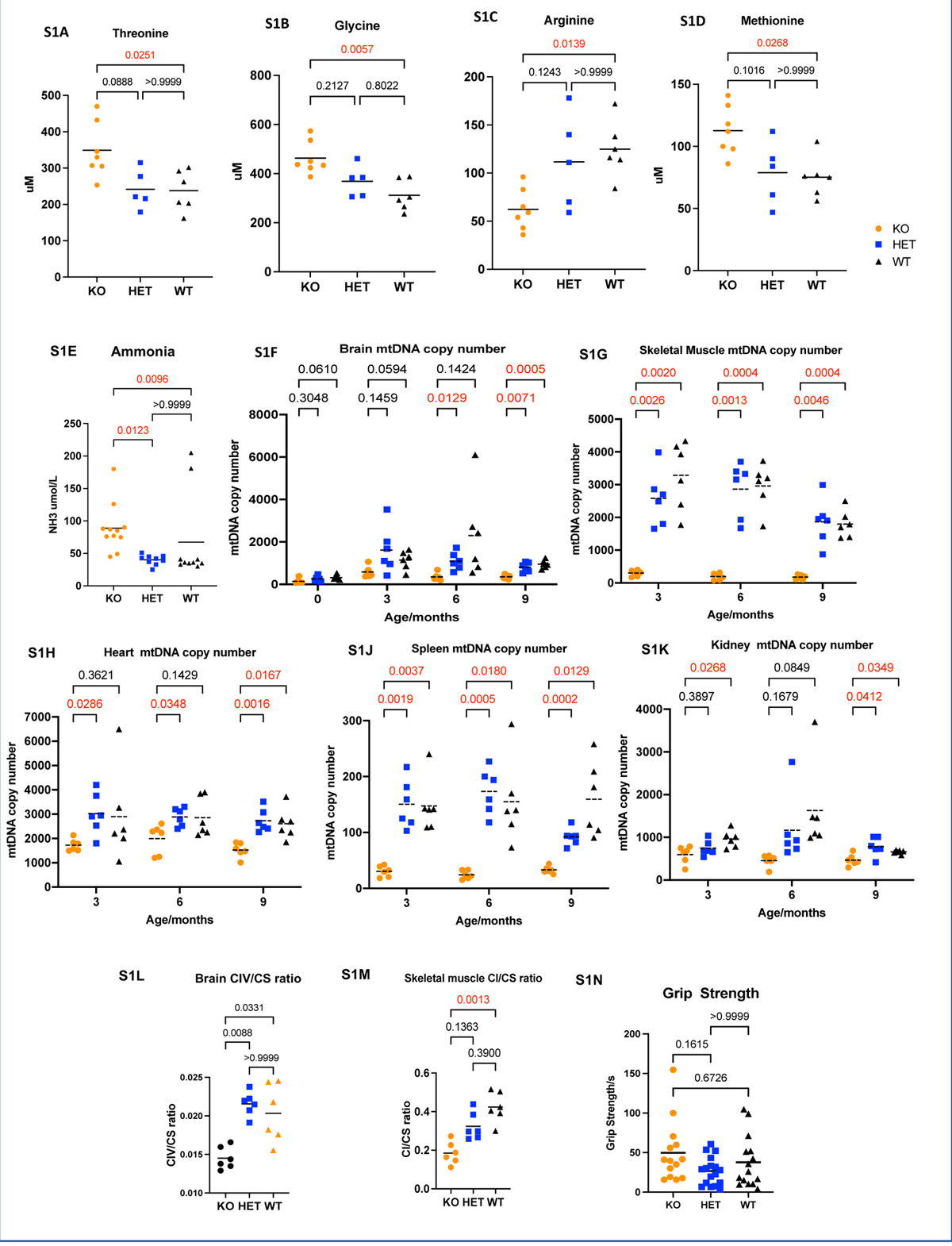
Supplementary baseline phenotyping data for Dguok KO mouse model **S1A-S1D:** Serum amino acid measurements for threonine, glycine, arginine and methionine. **S1E:** Hyperammonaemia was demonstrated in KOs. **S1F:** Brain mtDNA quantitation. KOs showed significant brain mtDNA depletion compared to WTs. **S1G, S1J:** Skeletal muscle and spleen mtDNA depletion is demonstrated from 3 months onward. **S1H, S1K** Heart and kidney mtDNA deficiency is demonstrated at 9 months **S1L:** KOs showed brain complex IV deficiency. **S1M:** KOs showed skeletal muscle complex I deficiency. **S1N:** Grip strength testing showed no differences between KOs and WT/HETs. Statistics: Kruskal-Wallis test

**Figure S2.**
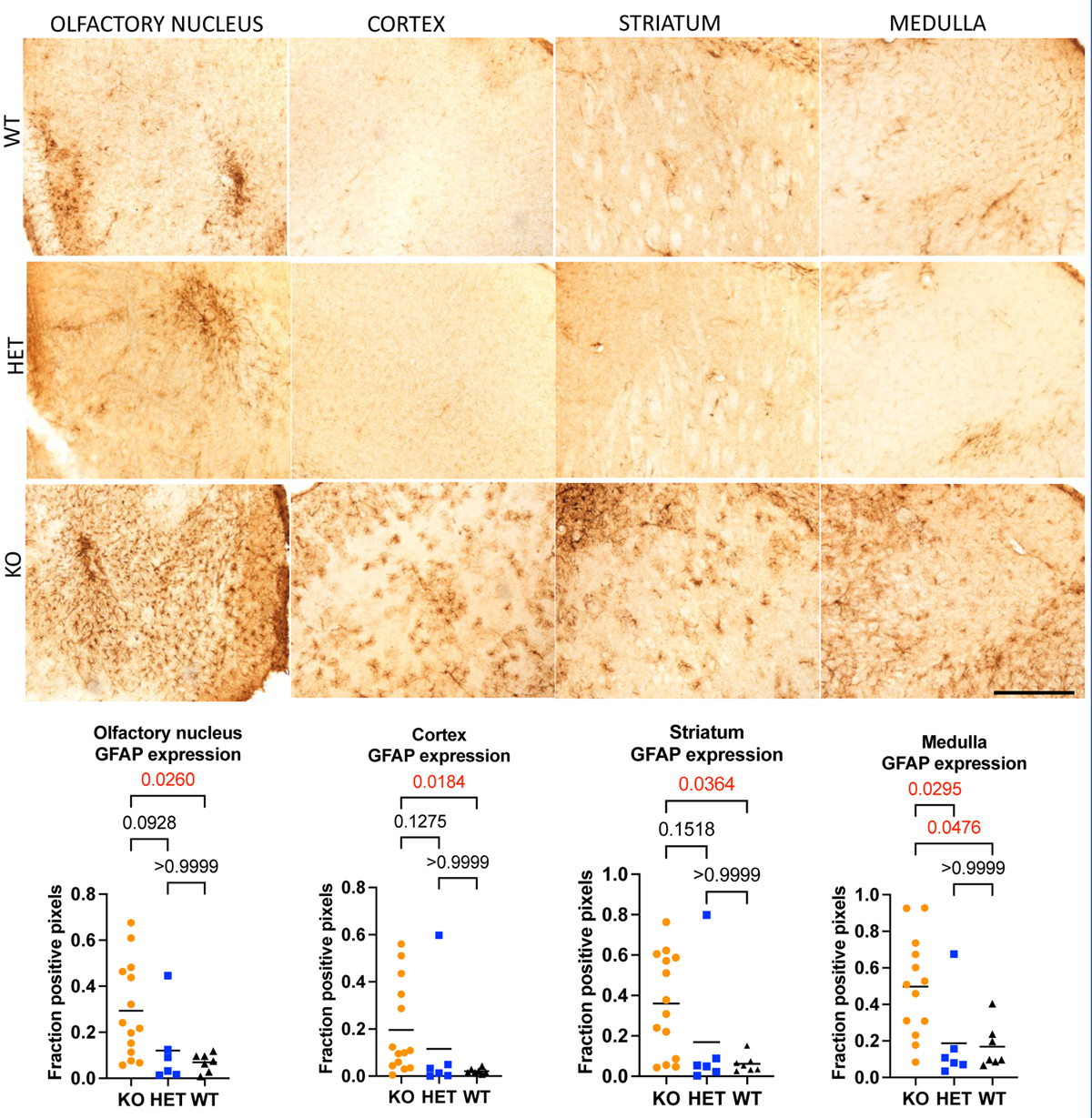
Characterisation of neurological phenotype in murine *Dguok* KO model. Neuroinflammatory phenotyping and detection of widespread neuroinflammation in *Dguok* KO mice was seen at nine months. Astrogliosis was evident on anti-GFAP immunohistochemistry in diffuse brain regions (olfactory nucleus, cortex, striatum and medulla oblongata) of KO mice in baseline phenotyping. Images were acquired at 10x magnification, scale bar: 500um.

**Figure S3.**
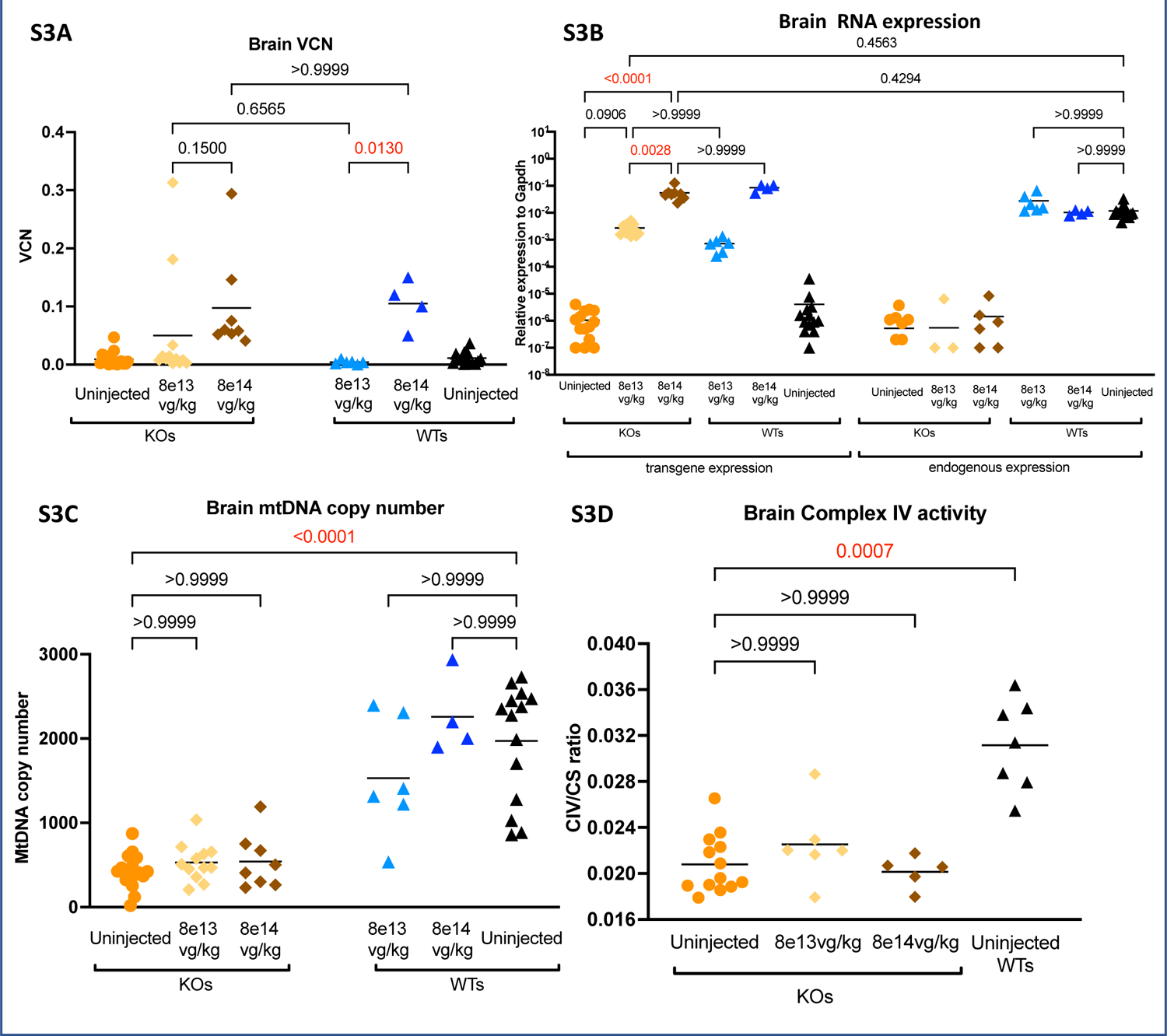
Brain outcomes following neonatal gene transfer **S3A** Brain VCN data in injected KO and WT mice showed low transduction **S3B** Brain RNA expression data in injected KO and WT mice**. S3C** Brain mtDNA copy number data in injected KO and WT mice**. S3D** Brain complex IV activity in injected KO mice. Statistics: Kruskal-Wallis test.

**Figure S4.**
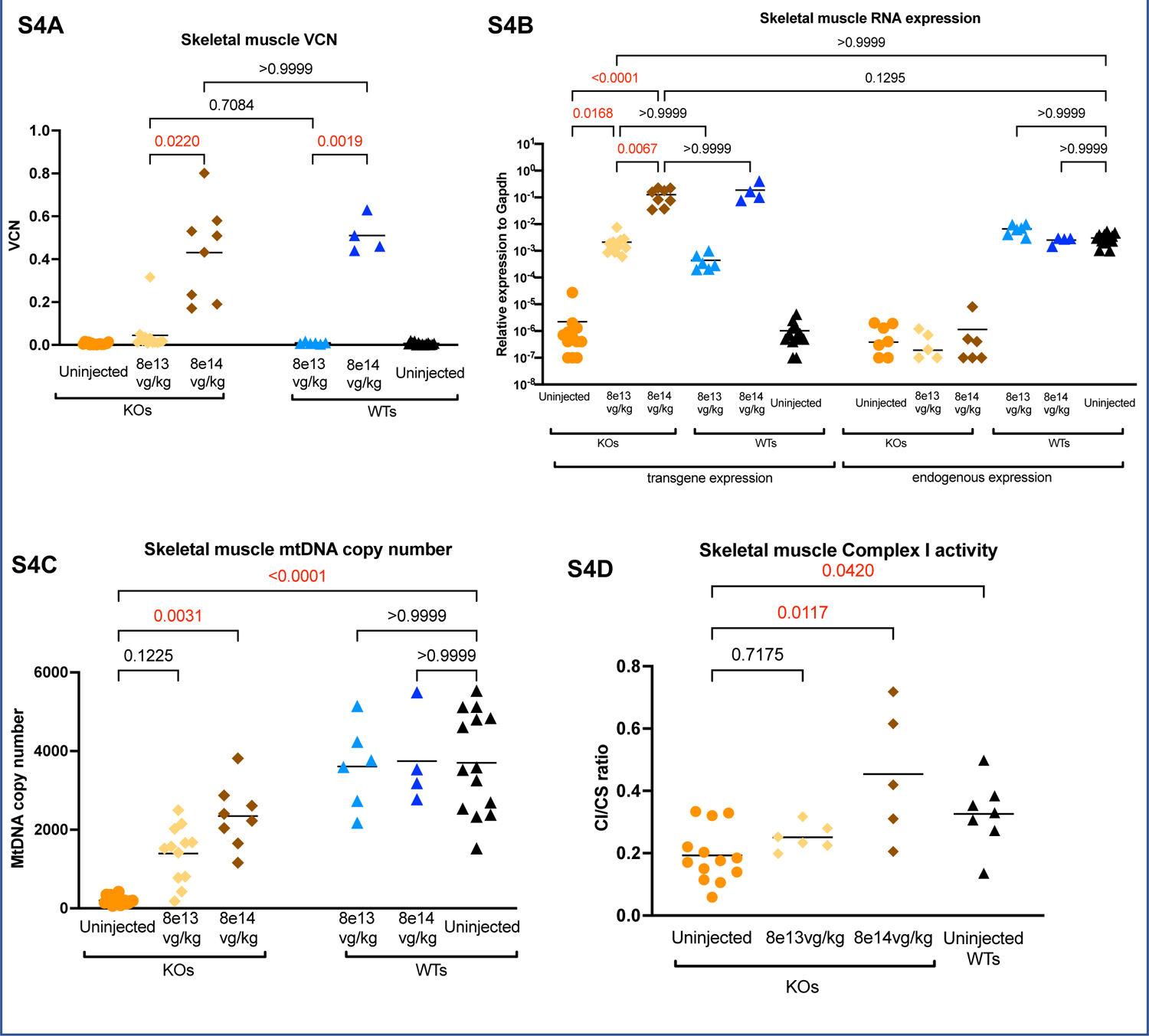
Outcomes of neonatal gene transfer in skeletal muscle. **S4A** Skeletal muscle VCN data in injected KO and WT mice. **S4B** Skeletal muscle RNA expression data in injected KO and WT mice**. S4C** Skeletal muscle mtDNA copy number data in injected KO and WT mice**. S4D** Skeletal muscle complex I activity in injected KO mice. Statistics: Kruskal-Wallis test.

**Figure S5.**
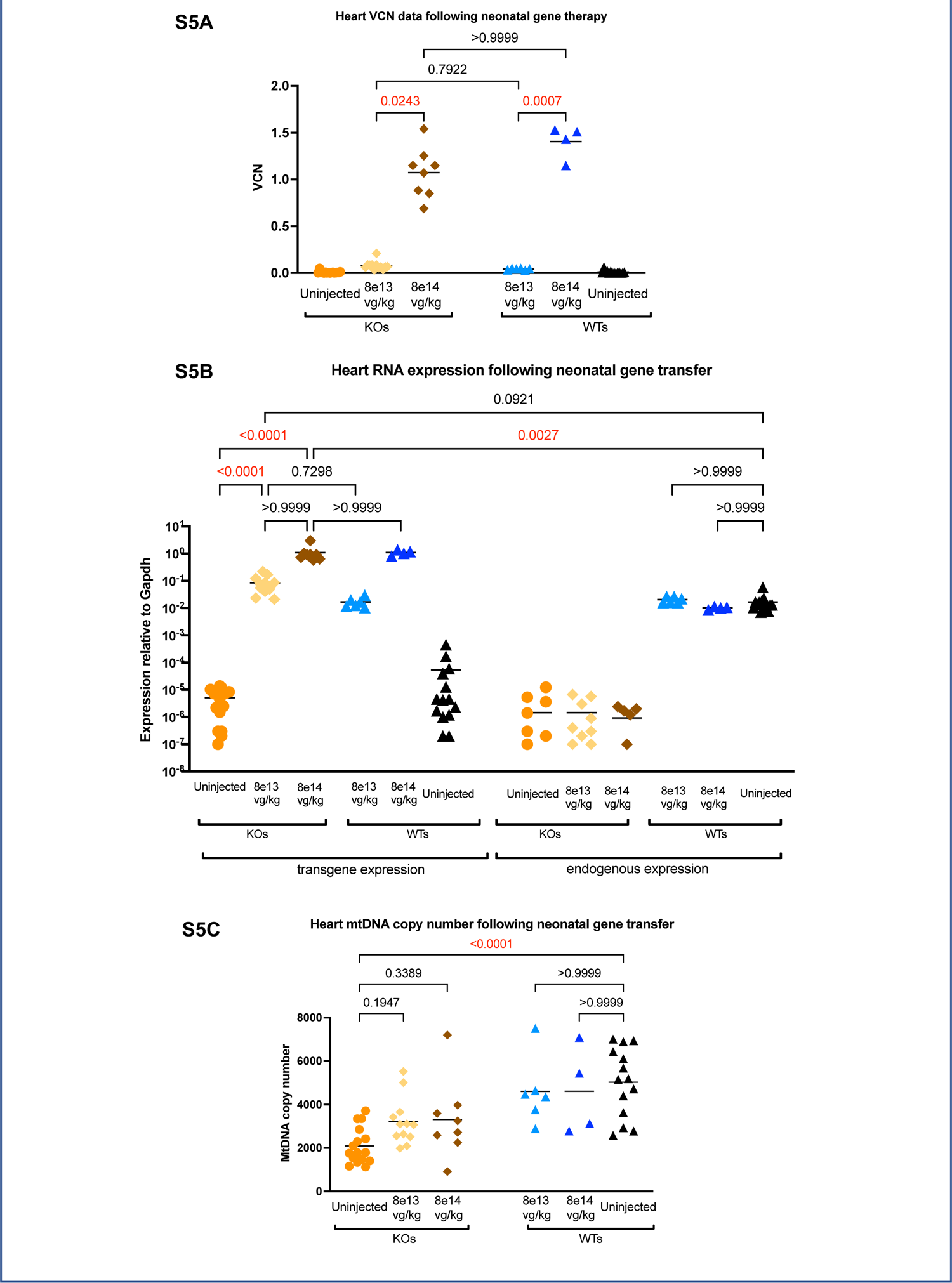
Outcomes of neonatal gene transfer in heart. **S5A** Cardiac VCN data in injected KO and WT mice. **S5B** Cardiac RNA expression data in injected KO and WT mice**. S5C** Cardiac mtDNA copy number data in injected KO and WT mice. Statistics: Kruskal-Wallis test.

---

**Supplementary Table 1.** OXPHOS measurements in baseline phenotyping. Blue: Significantly decreased activity in liver of KOs compared to WTs for complex I: p=0.0006, complex II+III p=0.038, complex III p=0.024, complex IV p=0.002). Red: Significantly increased citrate synthase activity in liver of KOs compared to WTs, p=0.008. Statistics performed using one-way ANOVA

|  |  | OXPHOS complex |  |  |  |  |
| --- | --- | --- | --- | --- | --- | --- |
| Organ |  | Complex I ratio | Complex II+III ratio | Complex III ratio | Complex IV ratio | Citrate synthase nmol/min/mg |
| Liver | KO | 0.348 | 0.151 | 0.034 | 0.023 | 62 |
|  | HET | 0.541 | 0.475 | 0.089 | 0.063 | 55 |
|  | WT | 0.789 | 0.424 | 0.084 | 0.083 | 41 |
| Brain | KO | 0.154 | 0.055 | 0.033 | 0.014 | 170 |
|  | HET | 0.218 | 0.052 | 0.058 | 0.021 | 178 |
|  | WT | 0.177 | 0.048 | 0.046 | 0.020 | 179 |
| Skeletal muscle | KO | 0.185 | 0.080 | 0.020 | 0.014 | 74 |
|  | HET | 0.324 | 0.144 | 0.034 | 0.019 | 70 |
|  | WT | 0.423 | 0.140 | 0.045 | 0.019 | 86 |
| Heart | KO | 0.529 | 0.034 | 0.009 | 0.007 | 397 |
|  | HET | 0.436 | 0.019 | 0.008 | 0.005 | 463 |
|  | WT | 0.451 | 0.027 | 0.006 | 0.005 | 463 |

## References

1. Viscomi C, et al. MtDNA-maintenance defects: syndromes and genes. J Inherit Metab Dis. 2017;40:587–599.

2. Moraes CT, et al. mtDNA depletion with variable tissue expression: a novel genetic abnormality in mitochondrial diseases. Am J Hum Genet. 1991;48:492–501.

3. Sezer T, et al. Novel deoxyguanosine kinase gene mutations in the hepatocerebral form of mitochondrial DNA depletion syndrome. J Child Neurol. 2015;30:124–8.

4. Johansson M, et al. Cloning and expression of human deoxyguanosine kinase cDNA. Proc Natl Acad Sci U S A. 1996;93:7258–62.

5. Saada A. Mitochondrial deoxyribonucleotide pools in deoxyguanosine kinase deficiency. Mol Genet Metab. 2008;95:169–73.

6. Labarthe F, et al. Clinical, biochemical and morphological features of hepatocerebral syndrome with mitochondrial DNA depletion due to deoxyguanosine kinase deficiency. J Hepatol. 2005;43:333–41.

7. Mandel H, et al. The deoxyguanosine kinase gene is mutated in individuals with depleted hepatocerebral mitochondrial DNA. Nat Genet. 2001;29:337–41.

8. Dimmock DP, et al. Clinical and molecular features of mitochondrial DNA depletion due to mutations in deoxyguanosine kinase. Hum Mutat. 2008;29:330–1.

9. Hanchard NA, et al. Deoxyguanosine kinase deficiency presenting as neonatal hemochromatosis. Mol Genet Metab. 2011;103:262–7.

10. Pronicka E, et al. Post mortem identification of deoxyguanosine kinase (DGUOK) gene mutations combined with impaired glucose homeostasis and iron overload features in four infants with severe progressive liver failure. J Appl Genet. 2011;52:61–6.

11. Kasapkara CS, et al. DGUOK-related mitochondrial DNA depletion syndrome in a child with an early diagnosis of glycogen storage disease. J Pediatr Gastroenterol Nutr. 2013;57:e28–9.

12. Al-Hussaini A, et al. Clinical and molecular characteristics of mitochondrial DNA depletion syndrome associated with neonatal cholestasis and liver failure. J Pediatr. 2014;164:553–9 e1-2.

13. Unal O, et al. Deoxyguanosine kinase deficiency: a report of four patients. J Pediatr Endocrinol Metab. 2017;30:697–702.

14. El-Hattab AW, et al. Deoxyguanosine Kinase Deficiency. In: M. P. Adam, H. H. Ardinger, R. A. Pagon, S. E. Wallace, L. J. H. Bean, K. Stephens and A. Amemiya, eds. GeneReviews((R)) Seattle (WA); 1993.

15. Dimmock DP, et al. Abnormal neurological features predict poor survival and should preclude liver transplantation in patients with deoxyguanosine kinase deficiency. Liver Transpl. 2008;14:1480–5.

16. Grabhorn E, et al. Long-term outcomes after liver transplantation for deoxyguanosine kinase deficiency: a single-center experience and a review of the literature. Liver Transpl. 2014;20:464–72.

17. Di Meo I, et al. Effective AAV-mediated gene therapy in a mouse model of ethylmalonic encephalopathy. EMBO Mol Med. 2012;4:1008–14.

18. Biousse V, et al. Long-Term Follow-Up After Unilateral Intravitreal Gene Therapy for Leber Hereditary Optic Neuropathy: The RESTORE Study. J Neuroophthalmol. 2021;41:309–315.

19. Bouaita A, et al. Downregulation of apoptosis-inducing factor in Harlequin mice induces progressive and severe optic atrophy which is durably prevented by AAV2-AIF1 gene therapy. Brain. 2012;135:35–52.

20. Cabrera-Perez R, et al. Alpha-1-Antitrypsin Promoter Improves the Efficacy of an Adeno-Associated Virus Vector for the Treatment of Mitochondrial Neurogastrointestinal Encephalomyopathy. Hum Gene Ther. 2019;30:985–998.

21. Di Meo I, et al. AAV9-based gene therapy partially ameliorates the clinical phenotype of a mouse model of Leigh syndrome. Gene Ther. 2017;24:661–667.

22. Lopez-Gomez C, et al. Synergistic Deoxynucleoside and Gene Therapies for Thymidine Kinase 2 Deficiency. Ann Neurol. 2021;90:640–652.

23. Bottani E, et al. AAV-mediated liver-specific MPV17 expression restores mtDNA levels and prevents diet-induced liver failure. Mol Ther. 2014;22:10–7.

24. Cunningham SC, et al. Gene Delivery to the Juvenile Mouse Liver Using AAV2/8 Vectors. Mol Ther. 2008;16:1081–1088.

25. Nathwani AC, et al. Long-term safety and efficacy following systemic administration of a self-complementary AAV vector encoding human FIX pseudotyped with serotype 5 and 8 capsid proteins. Mol Ther. 2011;19:876–85.

26. Dalwadi DA, et al. AAV integration in human hepatocytes. Mol Ther. 2021;29:2898–2909.

27. Strom SC, et al. Chimeric mice with humanized liver: tools for the study of drug metabolism, excretion, and toxicity. Methods Mol Biol. 2010;640:491–509.

28. Ignatenko O, et al. Loss of mtDNA activates astrocytes and leads to spongiotic encephalopathy. Nat Commun. 2018;9:70.

29. Chand D, et al. Hepatotoxicity following administration of onasemnogene abeparvovec (AVXS-101) for the treatment of spinal muscular atrophy. J Hepatol. 2021;74:560–566.

30. Binny CJ, et al. Vector systems for prenatal gene therapy: principles of adeno-associated virus vector design and production. Methods Mol Biol. 2012;891:109–31.

31. Zhou X, et al. Severe mtDNA depletion and dependency on catabolic lipid metabolism in DGUOK knockout mice. Hum Mol Genet. 2019;28:2874–2884.

32. Hargreaves P, et al. Diagnostic value of succinate ubiquinone reductase activity in the identification of patients with mitochondrial DNA depletion. J Inherit Metab Dis. 2002;25:7–16.

33. Shepherd D, et al. The kinetic properties of citrate synthase from rat liver mitochondria. Biochem J. 1969;114:597–610.

34. King TE. Preparations of succinate—cytochrome c reductase and the cytochrome b-c1 particle, and reconstitution of succinate-cytochrome c reductase. Methods in Enzymology. 1967;10:216–225.

35. Wharton DCT, A. Cytochrome oxidase from beef heart mitochondria. Methods in Enzymology. 1967;10:245–250.

36. C.I. Ragan MTW, V.M. Darley-Usmar, P.N. Lowe. In: D. R. V.M. Darley-Usmar, M.T. Wilson ed. Mitochondria: A Practical Approach Oxford, England IRL Press; 1987: 79–113.

37. Baruteau J, et al. Argininosuccinic aciduria fosters neuronal nitrosative stress reversed by Asl gene transfer. Nat Commun. 2018;9:3505.

38. Cohen SA, et al. PITC derivatives in amino acid analysis. Nature. 1986;320:769–70.

39. Pramanik BC, et al. Identification of phenylthiocarbamyl amino acids for compositional analysis by thermospray liquid chromatography/mass spectrometry. Anal Biochem. 1989;176:269–77.

